# Probing the physiological role of the plastid outer-envelope membrane using the oemiR plasmid collection

**DOI:** 10.1101/2023.07.20.549935

**Authors:** Serena Schwenkert, Wing Tung Lo, Beata Szulc, Chun Kwan Yip, Anna I. Pratt, Siobhan A. Cusack, Benjamin Brandt, Dario Leister, Hans-Henning Kunz

**Author notes:** Corresponding author: Serena Schwenkert, & Hans-Henning Kunz.

## Abstract

Plastids are the site of complex biochemical pathways, most prominently photosynthesis. The organelle evolved through endosymbiosis with a cyanobacterium, which is exemplified by the outer envelope (OE) membrane that harbors more than 40 proteins in Arabidopsis. Their evolutionary conservation indicates high significance for plant cell function. While a few proteins are well-studied as part of the protein translocon complex the majority of OE protein (OEP) functions is unclear. Gaining a deeper functional understanding has been complicated by the lack of observable loss-of-function mutant phenotypes, which is often rooted in functional genetic redundancy. Therefore, we designed OE-specific artificial micro RNAs (oemiRs) capable of downregulating transcripts from several loci simultaneously. We successfully tested oemiR function by performing a proof-of-concept screen for pale and cold-sensitive mutants. An in-depth analysis of pale mutant alleles deficient in the translocon component TOC75 using proteomics provided new insights into putative compensatory import pathways. The cold stress screen not only recapitulated three previously known phenotypes of cold-sensitive mutants, but also identified four mutants of additional oemiR OE loci. Altogether our study revealed a role of the OE to tolerate cold conditions and showcasts the power of the oemiR collection to research the significance of OEPs.

## Introduction

Plant chloroplasts are the cellular site of photosynthesis and host a number of interwoven pathways central to plant metabolism. Chloroplasts represent one specific type of plastid. However, all plastids originate from a single endosymbiotic event involving a cyanobacterial-like cell and an ancient host cell. As a result, plastids are enclosed by double-membranes consisting of the outer (OE) and inner (IE) envelope (Dyall *et al*., 2004).

Most studies have primarily focused on elucidating the IE function, as the OE was often regarded a nonspecific molecular sieve or as a remnant of the food vacuole from its engulfment during endosymbiosis (Day and Theg, 2018). Nevertheless, in recent years the significance of the OE and its proteins was demonstrated in various biological processes. The majority of OE proteins were in fact inherited from their prokaryotic ancestors, but key additions of eukaryotic descent required for integration of the engulfed cell are equally essential (Barth *et al*., 2022; Breuers *et al*., 2011). The mosaic nature of the membrane was instrumental for successful endosymbiosis and controls organelle biogenesis as well as plastid division.

In Arabidopsis, the OE harbors over 40 proteins (OEPs). OEPs thought to be of prokaryotic origin include porin-type channels, such as translocon of the outer chloroplast membrane (TOC)75, OEP21, OEP24, OEP37, and OEP40. Despite a lack of structural homology, they all function in preprotein and metabolite transport across the OE. Eukaryotic-type OEPs include tail-anchored proteins, such as OEP7 and OEP9, and GTPase receptors TOC34 and TOC159 (Barth *et al*., 2022). Apart from preprotein import and metabolite shuttling, OE proteins represent important players in other critical cellular functions, such as lipid biosynthesis (e.g., TGD4), or plastid division (PDV1/2) (Miyagishima *et al*., 2006; Xu *et al*., 2008). More recently, proteins belonging to the chloroplast-associated protein degradation (CHLORAD) pathway have unraveled an intriguing role of the OE in organellar protein ubiquitination and protein degradation (Ling *et al*., 2019; Woodson *et al*., 2015).

A plant’s ability to adjust to environmental perturbations depends heavily on changes in chloroplast metabolism. Consequentially, the import of nuclear encoded proteins as well as the shuttling of metabolites in and out of the chloroplast play a vital role in such acclimation processes (Kleine *et al*., 2021; Schwenkert *et al*., 2022). In recent years, proteins located in the chloroplast envelope including a number of OEPs have also been linked to plant acclimation responses in particular towards low temperature. In an extensive proteomics approach Trentmann et al., 2020 could show that several OEPs are differentially regulated after cold treatment.

Forward genetic screening for plant mutants with altered stress responses would be a powerful approach to identify these and additional OEPs with roles in acclimation *in planta*. Unfortunately, such screens have limited success rates when multiple genes encode for proteins with redundant functions (Cutler and McCourt, 2005). This hurdle can be overcome by using an artificial microRNA (amiR) approach with constructs that have the ability to target and downregulate multiple homologs that potentially serve similar functions (Hauser *et al*., 2013; Jover-Gil *et al*., 2014). In addition, amiR lines are mostly hypomorphic enabling the study of gene loss effects in loci which cause embryo lethality if lost entirely (Kunz *et al*., 2014b). Thus far, amiR screens have successfully helped to identify redundant proteins involved in processes such as auxin transport, abscisic acid signaling, arsenite and cadmium responses (Hauser *et al*., 2019; Xie *et al*., 2021; Zhang *et al*., 2018). Considering the fact that many OEPs are represented by gene families, amiR-based forward genetic screens provide an ideal approach to dissect the molecular fine-tuning capacities of this chloroplast protein subset.

In this study, we designed a collection of 36 binary pGreen-based vectors outfitted with OE-specific amiRNAs (oemiRs) targeting all to date verified OEPs. The tool was used to generate an initial oemiR plant mutant pool. As a proof of concept, this pool was screened for pale/photosynthesis-related as well as cold acclimation phenotypes. Since several OEPs are involved in preprotein import photosynthesis-related phenotypes pale plants were expected. To further investigate the aforementioned link between OEPs and cold acclimation we chose cold treatment as screening condition. One of the isolated pale plants was identified as a *toc75* loss-of-function mutant, a component of the protein import complex. This mutant was analyzed in more detail to understand the molecular consequences of impaired preprotein import on the cellular plant proteome.

## Methods and Material

### Genetic redundancy predictions

Genetic redundancy was predicted among gene pairs using the model described by (Cusack *et al*., 2021). Homologous gene pairs that clustered together in the phylogenetic tree were paired for analysis to determine the likelihood of their functional redundancy. Several genes that were related but did not cluster together in the phylogenetic analysis were also paired for model validation (Supplementary Data S1). Features were generated as described in the previous publication with one modification: coding sequences were here aligned in RAxML-NG using the JTT model (Cusack *et al*., 2021). The model implemented Random Forest and was trained on the “extreme redundancy” dataset with 200 features.

### oemiR plasmid collection generation and Agrobacterium trans/ormation

All target specific anti-sense and sense amiRs (amiR* and amiR, respectively) sequences were designed using the Web MicroRNA Designer, WMD3 (www.wmd3.weigelworld.org). Each amiR construct used in this study was cloned individually. The vector backbone was PCR amplified using the Platinum SuperFi II DNA Polymerase and the primers vec_fwd and vec_rev (Supplementary Data S1) (ThermoFisher Scientific), digested with DpnI and gel purified (Macherey&Nagel). The template for the vector backbone amplification was a fully functional and binary amiR expression clone in the pGREEN-based vector backbone vector called pG20_MCS_Hyg (Pratt *et al*., 2020). For each amiR construct, primers were designed with the amiR* and amiR being flanked by 5’ and 3’ sequences binding to the template vector (Supplementary Data S1). All amiR fragments were PCR amplified with the respective individual primer pairs using the Phusion polymerase (New England Biolabs) and subsequently gel purified (Macherey&Nagel). The vector backbone and the amiR fragments were assembled using Gibson seamless cloning (New England Biolabs) according to manufacturer’s instructions resulting on a functional binary expression vector. Maps of all vectors are provided in Supplementary Data S2.

### PCR analysis and sequencing

Genomic DNA was isolated according to Kotchoni and Gachomo (2009). Subsequently, the amiR construct was amplified from the genomic DNA with Taq-Polymerase PCR using the primers mir319_for and HSP18_rev. The PCR programme used consisted of an initial denaturation step of 30 seconds at 95°C. This was followed by a continued denaturation step of 30 seconds at 95°C, an annealing step for 50 seconds at 49°C and extension step for 1 minute at 68°C. This was repeated for a total of 36 cycles before a final extension step for 5 minutes at 68°C. PCR products were sequenced (Sanger sequencing, LMU Faculty of Biology, Genetics Sequencing Service).

### Plant growth and Arabidopsis trans/ormation

Arabidopsis plants were grown either on soil or on sterile solid ½ Murashige-Skoog-Medium (MS) medium. Plants were grown under long day conditions (day: 16 h 100 μmol photons m^-2^ s_-1_, 21°C; night: 8 h dark, 16°C) in a climate chamber (cold treatment and growth on plates) or the greenhouse. Treatment at 4°C was performed under the same light/dark regime. *Arabidopsis thaliana* Columbia (Col-0) accession was used as wild type strain. Initially, individual *Agrobacteria* cultures carrying one of the 36 plasmids each were grown as overnight cultures. The next morning, all cultures were normalized to the same OD_600_ and used to inoculate one joined 350 ml culture. Stably transformed Arabidopsis plants were generated using the floral dip method (Clough and Bent, 1998). The dipping procedure was repeated after one week. For selection of transformed plants, 1% agar containing ½ MS medium was supplemented with hygromycin. An ½MS mix with vitamins was used without the addition of sucrose. The medium was then adjusted to a pH of 5.7 with KOH before autoclaving and subsequent supplementation with 15 ug*ml^-1^ hygromycin. ln addition to non-selected Col-0 a transgenic line carrying a hygromycin resistance [control, (Kunz *et al*., 2014a)], was employed as a control. No apparent changes from Col-0 wild-type were observed.

### Chlorophyll (Chl) a /luorescence measurements

Chl *a* fluorescence of intact plants was measured using imaging pulse-amplitude-modulation fluorometry (Imaging PAM, Walz, Effeltrich, Germany) as described previously (Schneider *et al*., 2019). In brief, plants were dark-adapted for 20 min and exposed to a pulsed measuring light (intensity 1, gain 2, damping 1) and a saturating light flash (intensity 10) to calculate *F*_v_/*F*_m_, PSII quantum yield (ΦII = (Fm′ − F)/Fm′] and PS II regulated non-photochemical energy loss [ΦNPQ= (Fm − Fm′)/Fm′] and PSII quantum yield of non-regulated non-photochemical energy loss [ΦNO=F/Fm].

### Blue native (BN)-PAGE

Thylakoid membranes were were isolated and solubilized with 1% fs-dodecylmaltoside as described previously (Schwenkert *et al*., 2006). Samples were normalized according to fresh weight and separated on a NativePAGE™, 3 bis 12 % Bis-Tris gel (Invitrogen).

### Proteome analysis

Proteome analysis was performed using total protein extracts isolated from three-week old leaf material (four biological replicates). Protein preparation, trypsin digestion, and liquid chromatography-tandem mass spectrometry (LC-MS/MS) were performed as described previously (Marino *et al*., 2019). Raw files were processed using the MaxQuant software version 2.1.3.0 (Cox and Mann, 2008). Peak lists were searched against the Arabidopsis reference proteome (Uniprot, www.uniprot.org, version April 2021) using the built-in Andromeda search engine (Cox *et al*., 2011) with default settings and ‘mach-between-runs’ was enabled. Proteins were quantified using the label-free quantification algorithm (LFQ) (Cox *et al*., 2014). For hierarchical clustering the ANOVA significant differentially expressed proteins based on the log_2_ of the z-score normalized LFQ intensities. Envelope proteins were identified according to annotated envelope proteins in (Bouchnak *et al*., 2019), omitting ribosomal proteins.

### Computational analysis

CLC Main Workbench 20.4 (QIAGEN) was used to generate the phylogenetic tree. Downstream proteomics statistical analysis was performed using Perseus version 2.0.6.0 (Tyanova *et al*., 2016), R and RStudio. Enrichment analysis was performed with ShinyGO (http://bioinformatics.sdstate.edu/go) (Ge *et al*., 2020) and subcellular localization analysis was performed with SUBA5 (http://.suba.live) (Hooper *et al*., 2017).

## Results and Discussion

### Design o/ the oemiR plasmid collection

To design the oemiR plasmid collection we selected candidates with a high confidence in respect of their subcellular and OE localization, respectively. Almost all selected protein targets were confirmed in previous proteomics studies and/or by immunoblots or fluorescence-based methods. Table 1 provides either a representative reference for a proteomics dataset or a functional study investigating the localization. Since IE and OE cannot be properly separated in Arabidopsis, studies on *Pisum sativum* OE membranes identifying the respective paralogues were included (Simm *et al*., 2013). Moreover, key publications analyzing functions of the individual proteins are provided Table 1. For several OEPs distinct functions have been assigned in the last decades. These assignments range from the formation of membrane channels enabling protein and metabolite trafficking to plastid division, lipid biosynthesis and signaling. Since most functions are tied to their varying structural prerequisites we summarized structural classifications of all OE proteins in Table 1 (Fish *et al*., 2022). Interestingly, a number of OE proteins form fs-barrels, which are common in Gram-negative bacteria and mitochondrial membranes where they generally serve as porins. Among these, OEP23 and JASSY stand out, as their structure prediction suggest that they are incomplete fs-barrel proteins. JASSY belongs to the START/RHO_alpha_C/PlTP/Bet_v1/CoxG/CalC superfamily, that contains a conserved ligand-binding domain forming a large hydrophobic binding cavity, which gives rise to its function as an OPDA transporter (Guan *et al*., 2019; Radauer *et al*., 2008). OEP23 has been shown to possess ion permeability, however, the exact transport substrates remain to be identified (Goetze *et al*., 2015). Further studies are also required to analyze the structural details of these potential incomplete and unusual fs-barrel proteins.

**Table 1:**
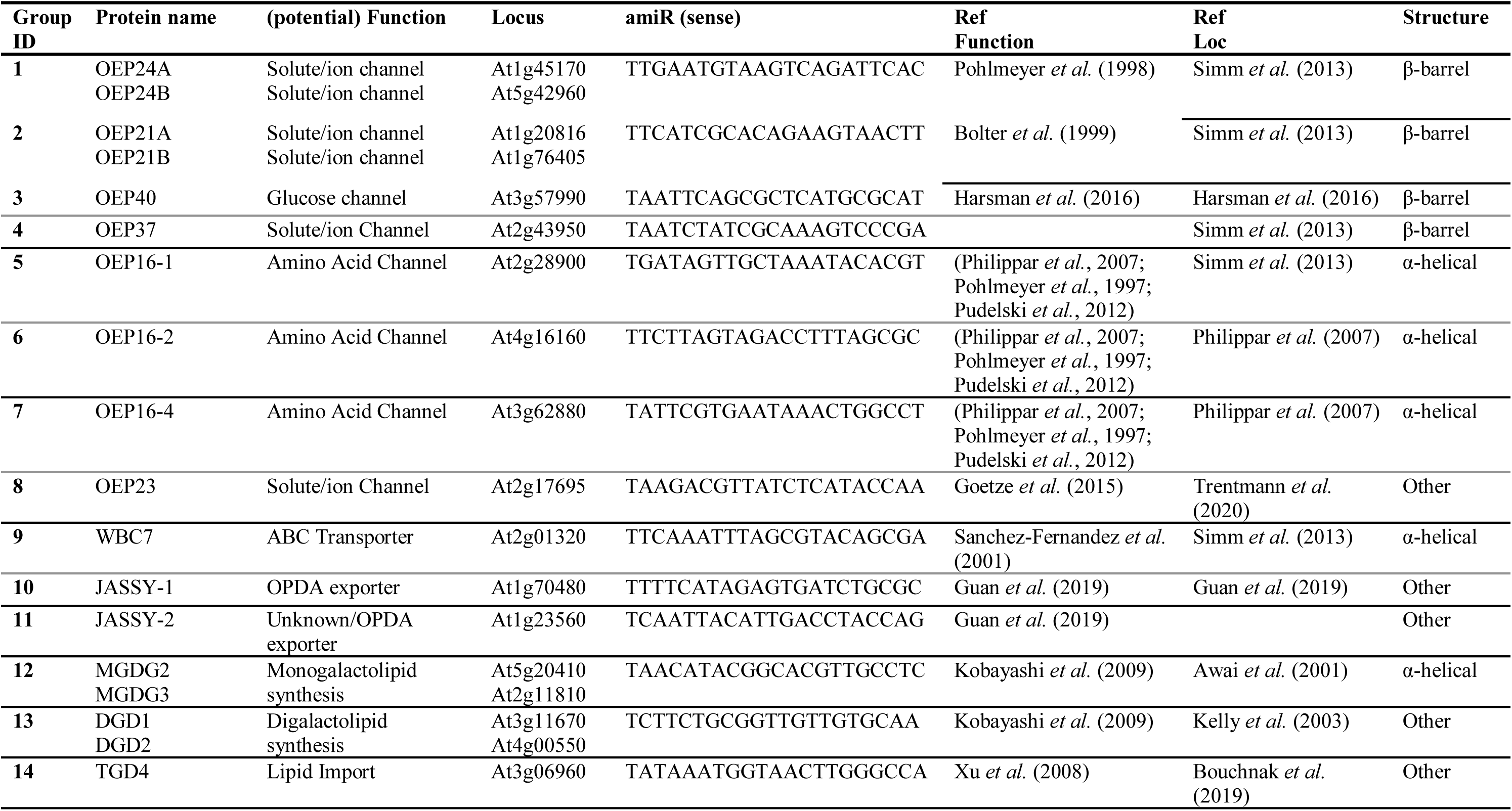
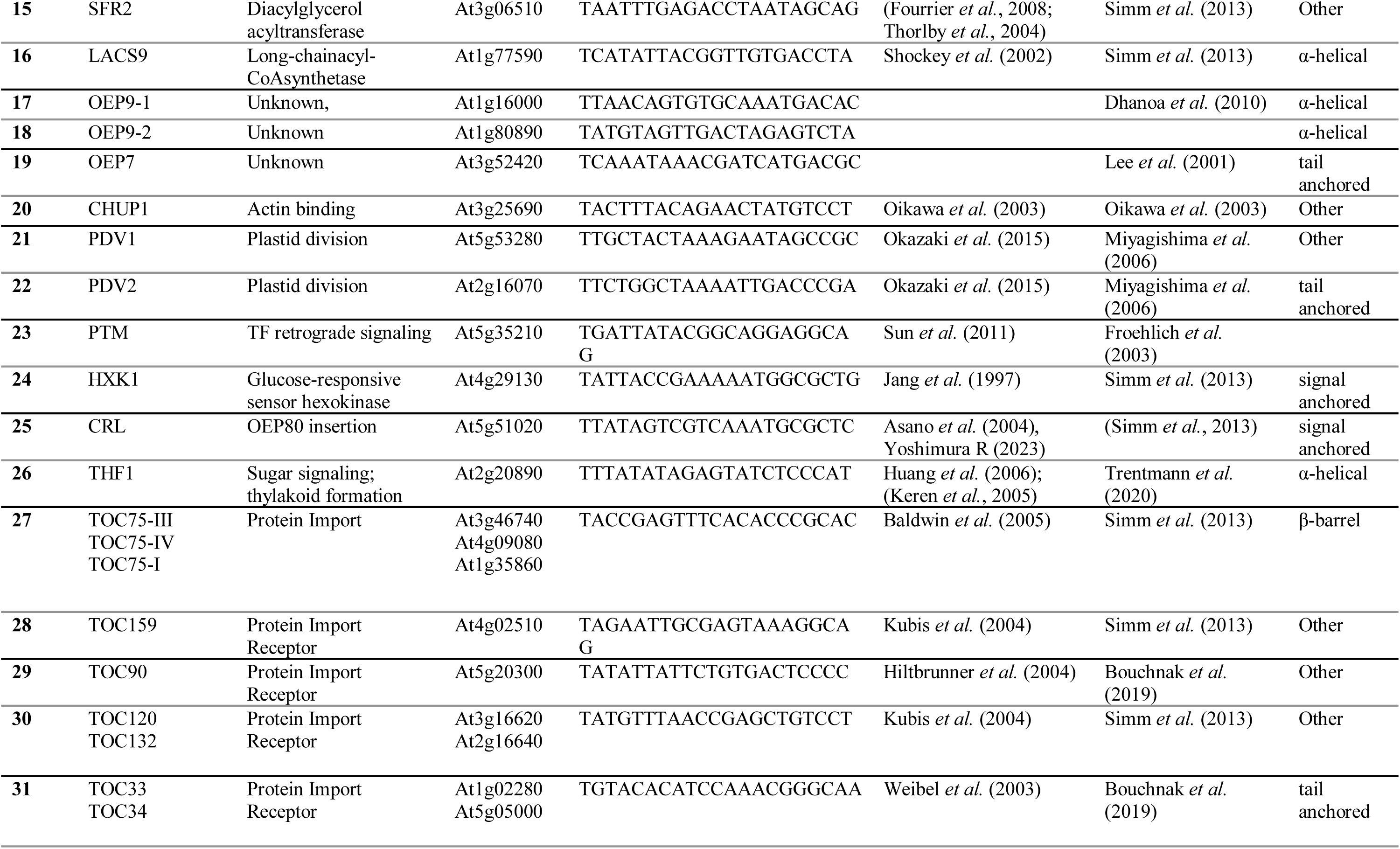
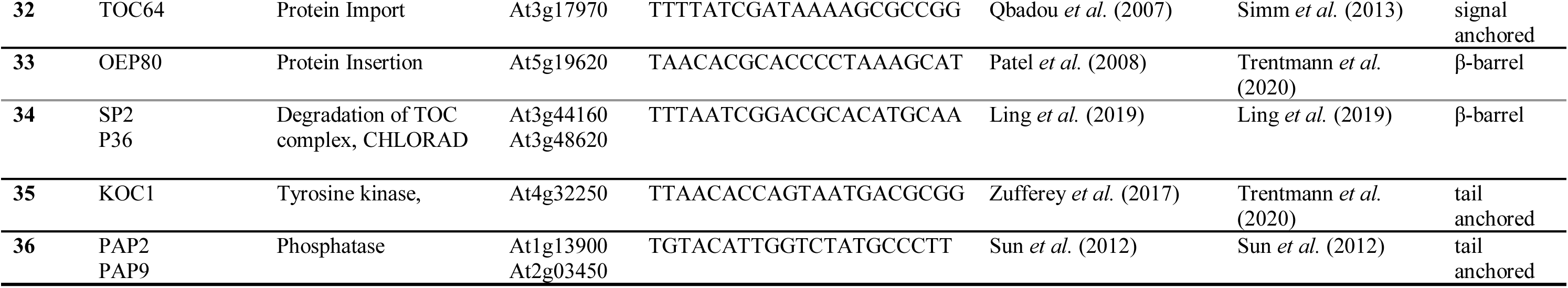
Targets of the oemiR plasmid collection.

To gain insight into the issue of functional redundancy among OE proteins, we aligned all cDNA sequences and compared these in a radial phylogenetic tree to identify gene clusters (Figure 1A). Several gene pairs or highly similar gene family members became apparent by clustering. Next up, the OE gene pairs were evaluated to determine their relative likelihood of functional redundancy, i.e., the risk that no phenotype would emerge in a single loss-of-function mutant. Indeed, the redundancy model suggested that there was a high risk of functional redundancy (Redundancy score ≥ 0.5) for all except one of the homologous gene pairs analyzed (Figure 1A and B).

**Figure 1:**
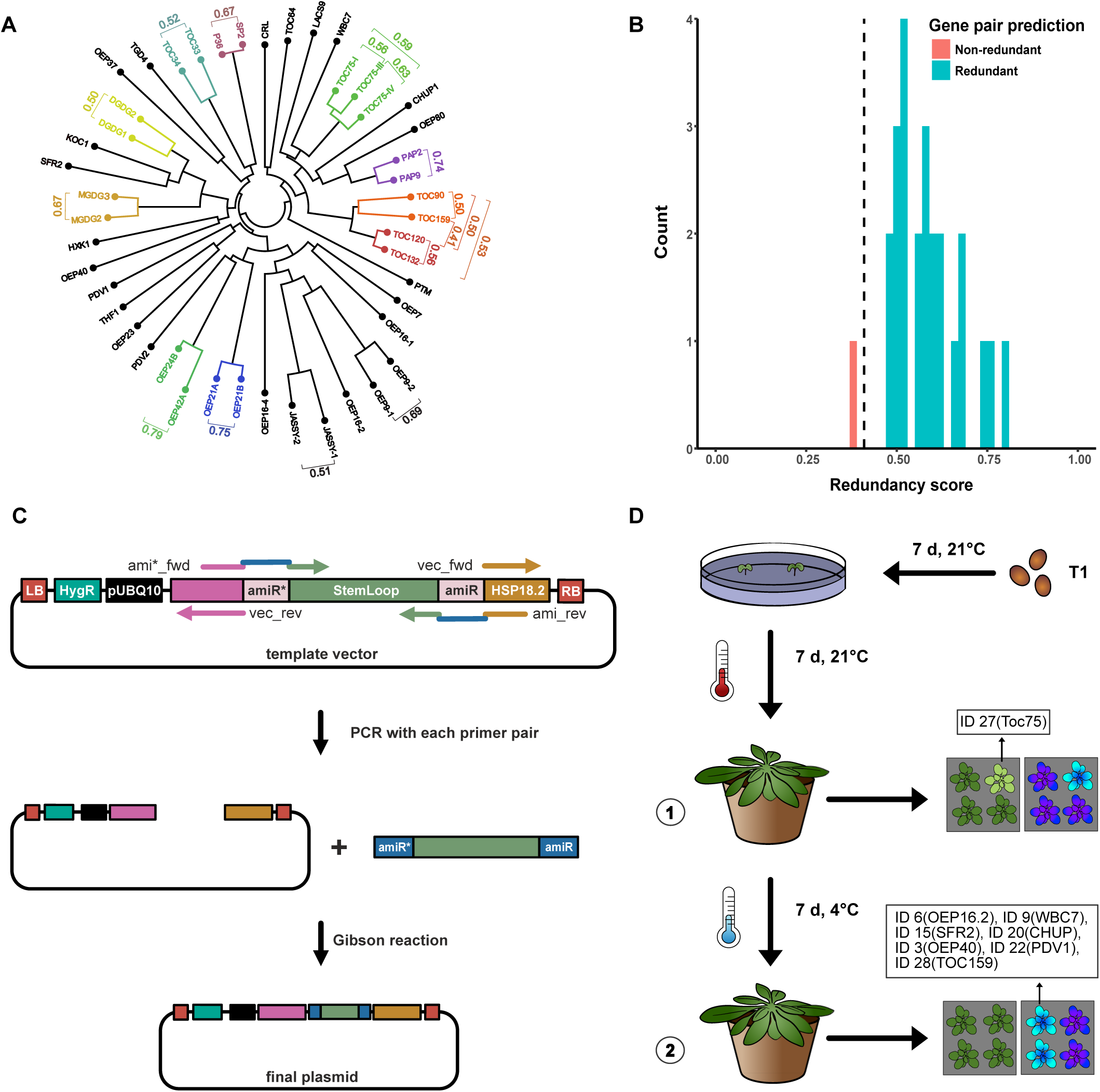
Design of the *oemiR* plasmid collection. **(A)** Circular phylogram based on all 46 OEP cDNA sequences. Colored genes indicate gene pairs targeted by one amiR. Scores adjacent to gene pairs indicate the predicted likelihood of redundancy between the two genes, with 0 indicating that the genes were likely non-redundant and 1 indicating that the genes were likely redundant. **(B)** Predicted redundancy scores for 25 pairs of genes among the OEPs. Genes that cluster together tend to have higher redundancy scores (≥0.5) than those that do not cluster together, consistent with the closer genetic relationships among clustered genes. **(C)** One-Step amiR cloning workflow. Each construct was cloned individually by PCR amplifying the vector backbone and the amiR insert followed by ln-Fusion assembly, resulting in a fully functional binary vector. pUBQ10: Ubiquitin10 promoter; amiR* and amiR: Specific amiR antisense and sense sequence, respectively; HygR: Hygromycin resistance cassette; LB and RB: Left and right T-DNA border, respectively. **(D)** Plant mutant screening. T_1_ seeds obtained after transformation with the *oemiR* plasmid collection were selected for hygromycin resistant seedlings. At the age of seven days, resistant seedlings (80 in total) were transferred to soil and grown for seven days at 21°C before being transferred to 4°C for seven days. At stages 1 and 2 phenotypes were monitored, Chl *a* fluorescence measurements were taken and plants differing from the control were subjected to PCR analysis and sequencing. AmiR group lDs were identified as indicated.

The main goal of the oemiR design was to find the minimal number of amiRs capable of downregulating as many gene targets as possible to bypass potential functional redundancy. Encouragingly, amiRs could be designed for all predicted functional redundant gene pairs, except JASSY and OEP9. Overall, 36 amiR constructs were sufficient to target all 46 genes (as indicated with colors in Figure 1A). All amiR group IDs targeting one or more gene loci transcripts along with the used amiR sequences are listed in Table 1. To streamline the molecular cloning, we established a new amiR one-step cloning process (Figure 1C). lnitially, the original MlR319a employed for the design of amiRs (Schwab *et al*., 2006), was inserted into an updated binary pGREEN-based vector called pG20_MCS_Hyg (Pratt *et al*., 2020). The high-copy vector then served as a PCR template to a) to incorporate the target specific amiR* and amiR sequences into a new DNA fragment also containing the stem loop and b) to amplify the vector backbone. Subsequently, both fragments were assembled using Gibson seamless cloning (New England Biolabs).

### Work/low o/ the oemiR proo/-o/-concept screen

All 36 oemiR constructs were transformed into Col-0 plants (T_0_) using standard *Agrobacteria-* mediated floral-dip. The resulting hygromycin-resistant T_1_ progeny was subjected to a two-step screening procedure for mutants impaired in growth, leaf paleness, photosynthesis and/or sensitivity to cold treatment (Figure 1D). In total, 80 T_1_ plants were analyzed. At an age of 14 days (long day 16h light /8h dark period conditions) phenotypes were visually inspected. Additionally, the plants’ photosynthetic capacity was evaluated by Pulse-Amplitude-Modulation (PAM) chlorophyll fluorometry (timepoint 1). *F*_v_/*F*_m_, the maximum quantum efficiency of photosystem ll (PSll), was evaluated as a general parameter reflecting plant fitness. Subsequently, plants were transferred to long day 16h/8h conditions at 4°C. After seven days, *F*_v_/*F*_m_ was measured again (timepoint 2). Generally, most plants did not show any morphological or other noticeable abnormalities. Nevertheless, some plants were smaller and paler than the control, already at timepoint 1. These plants were therefore subjected to PCR and sequencing to identify the causative amiR.

### Analysis o/ amiR-toc75 plants

Upon sequencing analysis of the pale plants described above, amiR ID 27, targeting the *TOC75* gene family, was identified. Since TOC75 is responsible for the import of most nuclear-encoded chloroplast proteins, this phenotype matched our expectations. Moreover, reduced import rates leading to a similar phenotype in a *toc75-III* RNAi mutant allele were observed previously (Huang *et al*., 2011). TOC75 proteins are part of the outer membrane protein of 85 kDa (Omp85) superfamily also found in the outer membranes of gram-negative bacteria and mitochondria (Day *et al*., 2019; Hsu and Inoue, 2009). Initially, three genes were assigned to the *TOC75* family. Apart from the ubiquitously expressed *TOC75-III, TOC75-IV* that might play a role in etioplasts is part of the family, as well as *TOC75-I*, which was classified as a pseudogene (Baldwin *et al*., 2005; Jackson-Constan and Keegstra, 2001). Later, TOC75-V/OEP80 was identified via proteomics studies as a TOC75 paralogue (Eckart *et al*., 2002) involved in the insertion of other OE β-barrel proteins (Gross *et al*., 2021; Patel *et al*., 2008). Complete loss of OEP80 renders mutants embryolethal (Patel *et al*., 2008). Since also TOC75-III null mutants are embryolethal (Baldwin *et al*., 2005), we chose the *amiR-toc75* lines to a) verify the specificity of our amiRNA constructs and b) to analyze how TOC75 downregulation affects the plant proteome in more detail.

To confirm the observed phenotype as caused by amiR ID 27, we performed an independent transformation with *Agrobacteria* only containing this plasmid. AmiR ID 27 was designed to target only *TOC75-I, TOC75-III* and *TOC75-IV*. Due to its distinct function in chloroplast biogenesis a separate amiR was designed for OEP80 (Table 1). Nevertheless, since the amiR ID 27 target site displayed some similarly to OEP80, this amiR represented an ideal tool to verify targeting specificity of our designed oemiRs (Figure 2A). Two independent *amiR-toc75* mutant lines were obtained and the progenitors from this transformation both yielded pale plant individuals in the T_1_ and T_2_ generations (Figure 2B). As a first step, a BN-PAGE was performed to investigate the overall integrity of thylakoid membrane complexes. All photosynthetic complexes were found to be reduced, especially PSll-LHCll supercomplexes, pointing towards a pleiotropic phenotype caused by lack of nuclear-encoded chloroplast proteins (Figure 2C).

**Figure 2:**
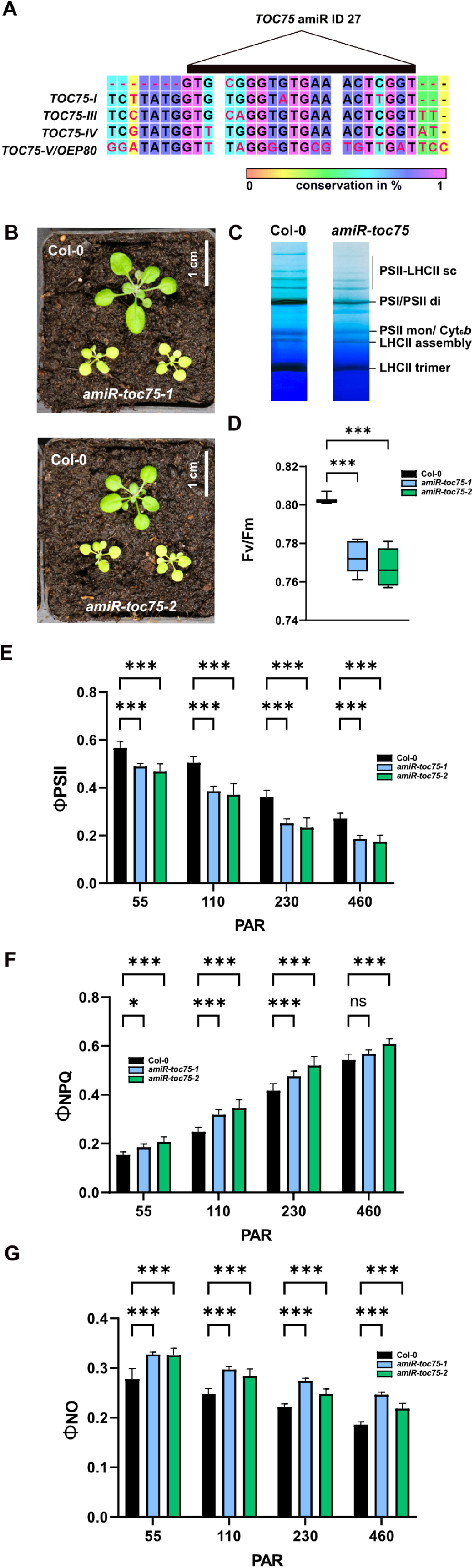
Analysis of *amiR-toc75* mutants. **(A)** Alignment of the amiR lD 27 target region. **(B)** Three-week-old *amiR-toc75* plants. **(C)** Large-pore BN gel of protein complexes from Col-0 and *amiR-toc75* chloroplasts. **(D-G)** Chl *a* fluorescence measurements of Col-0 and *amiR-toc75* plants. The asterisks indicate statistically significant differences according to one-way ANOVA comparing Col-0 to each mutant (*p<0.05; **p<0.01; ***p<0.001; n = 8).

To further analyze the photosynthetic performance of the mutants we measured chl *a* fluorescence parameters at increasing light intensities. *F*_v_/*F*_m_ was found slightly, but significantly reduced as compared to Col-0 wild-type plants (Figure 2D). Moreover, both *amiR-toc75* alleles displayed reduced PSII yield (ΦII), probably caused by a lower abundance of PSII complexes, which harbor a number of nuclear encoded subunits. This was accompanied by higher regulated non-photochemical quenching (ΦNPQ) and slightly elevated nonregulated non-photochemical quenching (ΦNO) indicative of oxidative stress cause by plastid malfunction in *amiR-toc75* plants (Figure 2E-G).

Next up, we were interested to analyze the extent of TOC75 protein downregulation by the amiR. Additionally, we also expected to gain a global understanding of the molecular consequences and potential compensatory mechanisms in response to loss of a protein import translocon unit. Therefore, we performed label-free protein quantification on leaf extracts using mass spectrometry (Figure 3A-F). Of the three predicted amiR targets (*TOC75-I*, *TOC75-III*, *TOC75-IV*), only TOC75-III protein was identified. This can be explained with the low TOC75-IV abundance in leaf chloroplasts (Baldwin *et al*., 2005). *TOC75-I* is a pseudogene as mentioned above. Nevertheless, TOC75-III was shown to be substantially downregulated by 3-fold in *amiR-toc75-1* and 3.6-fold in *amiR-toc75-2,* respectively. All other significantly differentially regulated proteins in both *amiR-toc75* vs. Col-0 were identified by performing a Student *t*-test. The -log_2_ fold changes of all up-and downregulated proteins were plotted against the *p*-value and depicted as a volcano plot (Figure 3A and Supplementary Data 3). Notably, OEP80 was unchanged in both mutant lines indicating a high target specificity towards TOC75-III by the amiR construct. In order to compare the two independently obtained mutant lines we matched the number of overlapping up-and downregulated proteins in both lines and found most proteins (between 62 - 75%) to be regulated in the same manner (Figure 3B).

**Figure 3:**
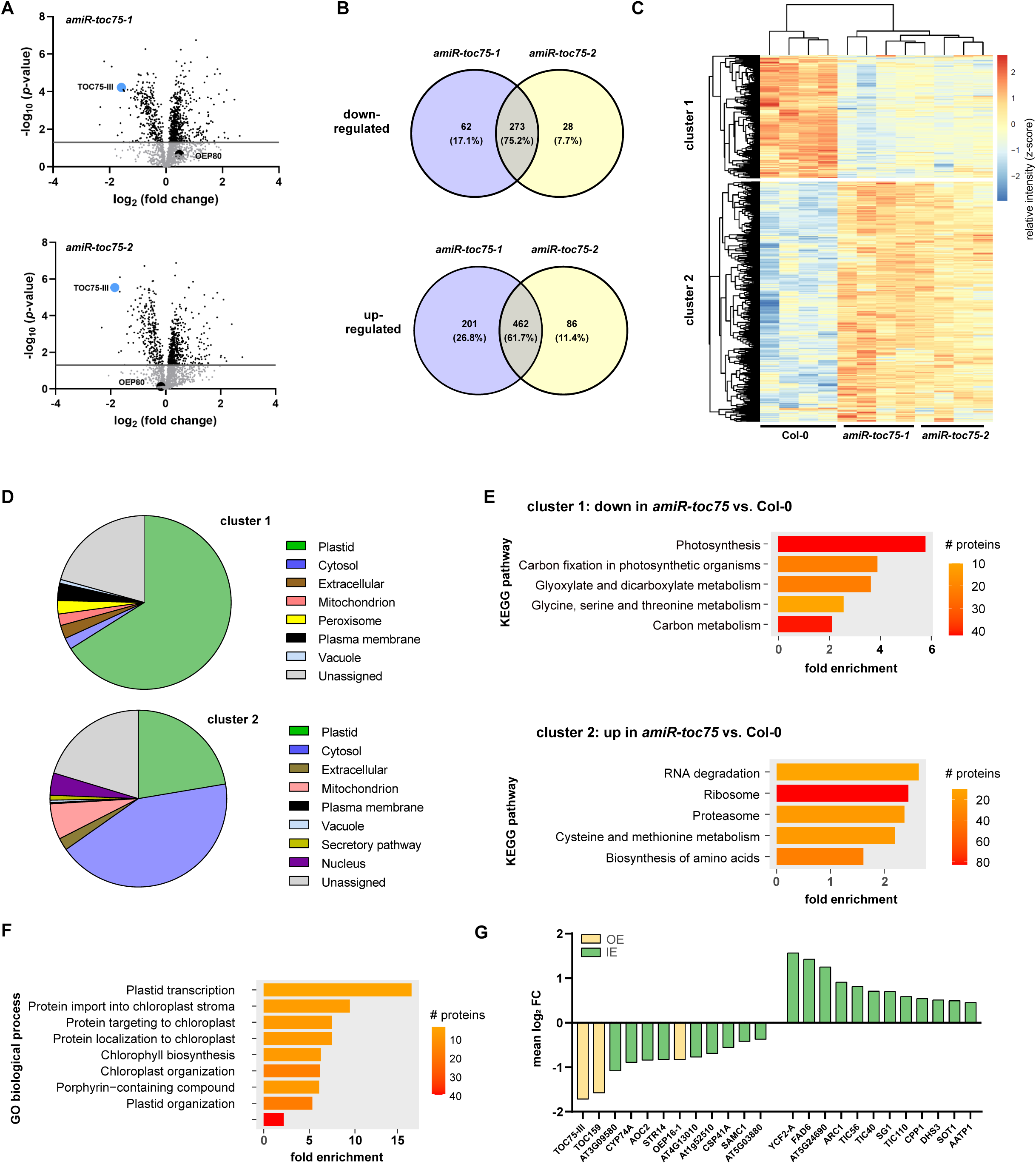
Proteome analysis of *amiR-toc75* mutants. **(A)** Volcano plots showing differentially regulated proteins in *amiR-toc75-l/amiR-toc75-2* vs. Col-0. Each dot represents one protein, plotted according to *p*-value and relative abundance ratio *amiR-toc75-l/amiR-toc75-2* and Col-0, upper and lower panel respectively. Toc75-lll and OEP80 are highlighted. The dashed line indicates the -log_10_ *p*-value of 1.5. **(B)** Venn diagrams showing the overlapping up-and downregulated proteins in both *amiR-toc75* lines. **(C)** Hierarchical cluster analysis of the differentially expressed proteins. Bar represents the relative z-score. **(D)** Subcellular enrichment analysis of the clusters identified in (C). **(E)** KEGG pathway enrichment analysis of the clusters identified in (C). **(F)** More than 2-fold up-and downregulated OE and lE proteins extracted from the clusters identified in (C). **(G)** GO biological process enrichment analysis of upregulated plastid proteins identified in (D).

Since not much is known about the effect of TOC75 downregulation on the composition of the proteome and in order to perform a functional enrichment analysis we subjected the data to hierarchical clustering analysis. Differentially regulated proteins grouped into two clusters. Cluster 1 contained 321 proteins, which were mainly downregulated in both *amiR-toc75* lines vs. Col-0. Cluster 2 held 619 mainly upregulated proteins in both *amiR-toc75* lines vs. Col-0 (Figure 3C). To estimate the compartmental protein abundance of down-vs. upregulated proteins we utilized the SUBA5 database. The results shown in Figure 3D reveal that the majority of downregulated proteins are localized in chloroplasts, which underpins the role of TOC75 in plastid preprotein import. Moreover, other components of the TOC complex, such as TOC159, a major preprotein receptor, were also downregulated in response to low TOC75 abundance (Figure 3E). Most of the upregulated proteins, however, were cytosolic. For further functional analysis of these proteins, we performed KEGG enrichment analysis. While cluster 1 was heavily enriched in proteins related to photosynthesis and chloroplast functions, the upregulated proteins (cluster 2) were enriched in proteins related to cytosolic ribosome assembly, translation and protein degradation (Figure 3F). One explanation is that compensatory mechanisms to level out the effects of reduced plastid protein import become initiated. This is achieved through higher translation rates and faster degradation of unimported, aggregated preproteins.

Interestingly, also a number of chloroplast upregulated proteins in *amiR-toc75* vs. Col-0 were identified. GO term enrichment analysis revealed that these comprised proteins with biological functions related to plastid transcription, but also proteins involved in preprotein import (Figure 3F). Among those were prominent proteins of the inner preprotein translocon (TIC) complex, i.e. TlC40 and TlC56. Also, TIC110 was upregulated, albeit it has been controversially discussed whether TIC110 is a permanent component of the TIC complex (Figure 3G) (Bolter, 2018; Jin *et al*., 2022; Nakai, 2015). In addition, Chaperonin CPN60, 93-kD heat shock protein HSP93-III/CLPC2 and chloroplast heat shock protein HSP90C are upregulated, which are also suggested to play a role in preprotein import (Bolter, 2018) (Supplementary Data S3). At first glance, an upregulation of these components seems to be a logic consequence if the cell tries to compensate reduced import rates across the OE. Nevertheless, it also bares the question how these proteins enter the chloroplast in the absence of a fully assembled functional TOC complex in the first place? Moreover, ten additional stromal proteins were upregulated more than two-fold. One possibility is that these proteins are preferentially recognized and transported to keep the mutant chloroplasts functional to some degree. Alternatively, some of these proteins might utilize non-canonical import pathways which are yet to be unraveled but have been posited in the past (Armbruster *et al*., 2009). In summary, this experiment confirmed that amiRs provide a reliable method to study loss-of-function effects in the OE circumventing functional genetic redundancy.

### ldenti/ication o/ mutants with lower photosynthetic per/ormance in the cold

A recent study has shown a significant adjustment of the OE proteome in response to cold treatments (Trentmann *et al*., 2020). To investigate the impact of OEP reduction on cold acclimation, we screened our initial *oemiR* mutant pool for phenotypic changes in response to low temperature. All 80 T_1_ plants were shifted into long-day growth conditions at 4°C. After one week at 4°C *F*_v_/*F*_m_ was determined. While the control and most *oemiR* mutants exhibited an *F*_v_/*F*_m_ value around 0.70, ten *oemiR* lines revealed average *F*_v_/*F*_m_ values of 0.55 and below (*p*<0.05 Student *t*-test). Subsequently, mutant plants with a cold-treatment induced drop in average *F*_v_/*F*_m_ were PCR genotyped and sequenced to identify the respective causative amiRs (Figure 4A and B). Out of these, seven mutant plants carried amiR constructs targeting *SENSlTlVE TO FREEZlNG2* (*SFR2)*, *CHLOROPLAST UNUSUAL POSlTlONlNG PROTElN l(CHUPl)*, *OEP40*, *TOCl59*, *OEPl6-2*, *PLASTlD DlVlSlON2* (*PDV2)*, and the putative ABC-type transporter *WBC7*, respectively. Three *oemiR* lines gave inconclusive sequencing results, potentially due to the presence of multiple insertions. In general, mutant lines carrying multiple amiRs can be cleaned up and turned into *oemiR* single mutant by backcrossing into Col-0 wild-type plants. The resulting F1 progeny needs to be tested for the heritability of the desired phenotype and genotyped.

**Figure 4:**
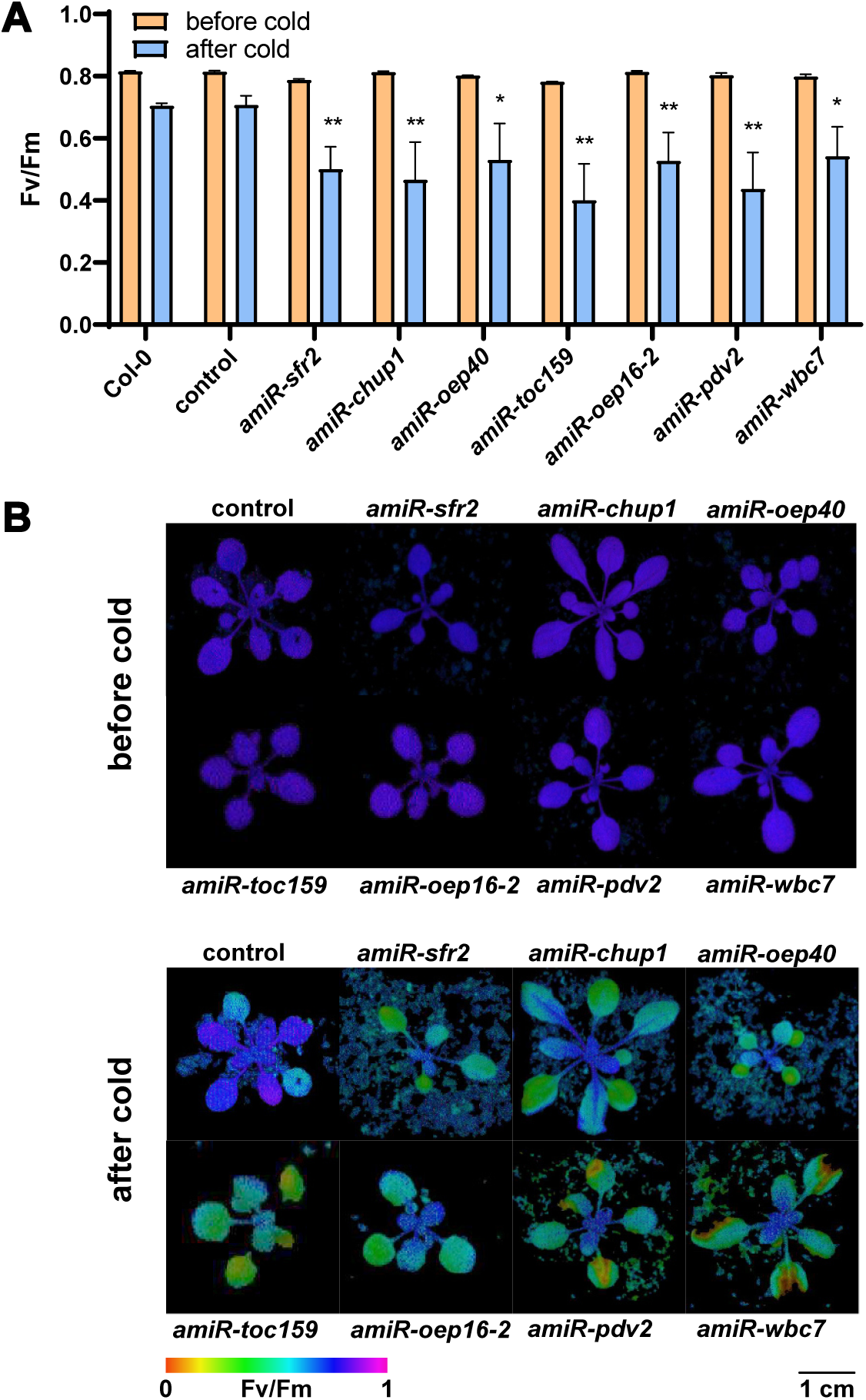
Screening of *oemiR* mutants with affected cold acclimation based on *F*_v_/*F*_m_ changes from wild-type control plants. **(A)** All mutant plants were treated at 4°C, with Chl *a* fluorescence measurements taken before and after treatment. Mutants that had an average *F*_v_/*F*_m_ value of 0.55 after cold treatment were identified and a Student *t*-test was performed after cold treatment with each mutant against the control (*p<0.05; **p<0.01; ***p<0.001; n = 4). Genotyping was performed on mutants that showed a statistically significant decrease in average *F*_v_/*F*_m_ after cold treatment (p < 0.05). Seven of these mutants were sequenced and identified to harbor amiR constructs targeting *SFR2, CHUPl, OEP40, TOCl59, OEPl6.2, PDV2* and *WBC7* respectively. **(B)** Chlorophyll fluorescence imaging before and after cold treatment.

A literature research confirmed that loss of functions in *s/r2*, *oep40* and *chup1* genes have been previously linked to cold-sensitivity. SFR2, a member of the Family-1 β-glycosidases, is partially responsible for removing monogalactolipids from the OE, a lipid-remodeling process vital for freezing tolerance (Moellering *et al*., 2010; Thorlby *et al*., 2004). Consequently, *s/r2* loss-of-function mutants suffer chloroplast damage after exposure to freezing conditions (Fourrier *et al*., 2008). Extended cold-treatments i.e., above 0°C, have not been reported thus far. *OEP40* encodes for a regulated β-barrel OE channel permeable to glucose and its phosphorylated derivatives. Loss of OEP40 triggers early flowering under cold temperature conditions, which is indicative of carbohydrate imbalance particularly in the floral meristem. (Harsman *et al*., 2016). Lastly, CHUP1 is an OEP featuring an actin binding domain responsible for chloroplast positioning and movement within leaf cells (Oikawa *et al*., 2003; Oikawa *et al*., 2008). Under ambient light/low temperature conditions *chup1* loss-of-function mutants exhibit lower *F*_v_/*F*_m_ values compared to control plants (Kitashova *et al*., 2021). This phenotype was recapitulated in the *amiR-chup1* line isolated in this study.

While the identification *SFR2*, *OEP40* and *CUP1* confirmed the functionality of our screening procedure, the three other cold-affected mutants identified are valuable for further studies directed towards the OE role in cold acclimation. The isolated mutants harbored individual amiR constructs targeting *TOCl59*, *OEPl6-2*, *PDV2*, and *WBC7* respectively. *TOC159*, also known as *PLASTlD PROTElN lMPORT 2* (*PPl2*), encodes for a membrane GTPase that functions as a transit-sequence receptor and represents an integral part required of the translocase complex. Previous work on *ppi2* mutants has shown that null alleles are seedling lethal but exhibit an albino phenotype if grown on sucrose (Bauer *et al*., 2000). The *amiR-tocl59* line isolated and used in this study did not show such a strong phenotype under normal growth conditions. Nevertheless, the mutant did exhibit cold-sensitivity. This finding underpins the value of milder mutant versions to study complex physiological stress responses. We did also find strongly comprised *amiR-toc159* lines reminiscent of the *ppi2* allele which we decided to not use for the cold sensitivity experiment.

OEP16-2 was shown to affect metabolic fluxes during ABA-controlled seed development and germination (Pudelski *et al*., 2012). It forms a potential amino-acid selective channel that possesses moderate cation-selectivity but high-conductance. While OEP16-1, which according to the design algorithm is not targeted by the *OEP16-2* amiR, is highly abundant in leaf tissue, the OEP16-2 isoform is preferably expressed during germination and late seed development (Pudelski *et al*., 2012). Nevertheless, the observed leaf phenotype in the *amiR-oep16-2* allele may indicate either unpredicted off-targeting towards OEP16-1 or a specialized role of OEP16-2 during cold acclimation which has previously not been reported. The same is true for PDV2, a tail-anchored protein that, along with its homolog PDV1, is a vital component of the division machinery in plastids (Chang *et al*., 2017; Miyagishima *et al*., 2006). Finally, WBC7 is a putative ABC transporter located in the OE (Sanchez-Fernandez *et al*., 2001). The exact molecular function of this protein remains elusive. Nevertheless, confirming our *amiR-wpc7* mutant phenotype results, WBC7 has been found to be down-regulated under cold conditions (Trentmann *et al*., 2020).

With this pilot screen we were able to emphasize a global function of OEPs in cold acclimation. However, to pursue the molecular functionalities causing the observed cold-sensitive phenotypes in *amiR-toc159*, *amiR-oep16-2*, *amiR-pdv2* and *amiR-wbc7* further verification steps are required. As exemplified for *amiR-toc75* individual oemiR constructs can be utilized to generate independent, yet comparable mutant lines with high specificity towards the gene(s) of interest. Homozygous T-DNA or CrispR mutants, if viable, can be employed to verify observed phenotype(s) and to unravel the biological role of the respective proteins in cold acclimation and beyond. Future studies for instance exploiting the *oemiR* collection will dissect the OE’s critical role for acclimation under other abiotic and biotic stress scenarios.

Overall, this OE-specific amiR-based screening tool exhibts great potential in overcoming common issues faced with others screening methods, such as seedling lethality and functional redundancy. The *oemiR* collection demonstrates the power of amiR-based strategies for new studies not limited to the plastid OE, but also other biological subsystems such as whole organelles.

## Supplementary data

The following supplementary data are available at JXB online.

**Supplementary Data St:** Primers used for cloning the oemiR plasmid collection and redundancy predictions for OEP gene pairs.

**Supplementary Data S2**: Vector maps of the oemiR plasmid collection.

**Supplementary Data S3:** Proteome analysis of *amiR-toc75* mutants.

## Acknowledgments

We thank Dr. Philip Day (WSU) for generating amiR-toc75 mutants and help with the initial design of the plasmid collection. We acknowledge excellent technical assistance by Yulia Davydova (LMU Munich), Denise Winkler (LMU Munich), and Alyssa Akamine (WSU). We further thank Susanne Muhlbauer (LMU Munich) for contributing and designing illustrations shown in Figure 1.

## Author contributions

SS, HHK: conceptualization and supervision; SS, WTL, BS, CKY, SAC, AIP, BB: investigation; DL: review and editing; SS, HHK, WTL: writing.

## Data availability

The mass spectrometry proteomics data have been deposited to the ProteomeXchange Consortium via the PRIDE partner repository with the dataset identifier PXD041299. Sequence information on the oemiR plasmid collection in provided in Supplementary Data S2. Plasmids are deposited at BCCM (https://bccm.belspo.be/deposit/public/plasmids).

